# Ivermectin repurposing for COVID-19 therapy: Safety and pharmacokinetic assessment of a novel nasal spray formulation in a pig model

**DOI:** 10.1101/2020.10.23.352831

**Authors:** J. Errecalde, A. Lifschitz, G. Vecchioli, L. Ceballos, F. Errecalde, M. Ballent, G. Marín, M. Daniele, E. Turic, E. Spitzer, F. Toneguzzo, S. Gold, A. Krolewiecki, L. Alvarez, C. Lanusse

**Author notes:** J. Errecalde and A. Lifschitz equally contributed to this work.

## Abstract

High ivermectin (IVM) concentrations suppress *in vitro* SARS-CoV-2 replication. Nasal IVM spray (N-IVM-spray) administration may contribute to attaining high drug concentrations in nasopharyngeal (NP) tissue, a primary site of virus entrance/replication. The safety and pharmacokinetic performance of a new N-IVM spray formulation in a piglet model were assessed. Crossbred piglets (10–12 kg) were treated with either one or two (12 h apart) doses of N-IVM-spray (2 mg, 1 puff/nostril) or orally (0.2 mg/kg). The overall safety of N-IVM-spray was assessed (clinical, haematological, serum biochemical determinations), and histopathology evaluation of the application site tissues performed. The IVM concentration profiles measured in plasma and respiratory tract tissues (nasopharynx and lungs) after the nasal spray treatment (one and two applications) were compared with those achieved after the oral administration. Animals tolerated well the novel N–IVM-spray formulation. No local/systemic adverse events were observed. After nasal administration, the highest IVM concentrations were measured in NP and lung tissues. Significant increases in IVM concentration profiles in both NP-tissue and lungs were observed after the 2-dose nasal administrations. The nasal/oral IVM concentration ratios in NP and lung tissues (at 6 h post-dose) markedely increased by repeating the spray application. The fast attainment of high and persistent IVM concentrations in NP tissue is the main advantage of the nasal over the oral route. These original results are encouraging to support the undertaking of further clinical trials to evaluate the safety/efficacy of the nasal IVM spray application in the treatment and/or prevention of COVID-19.

## Introduction

Ivermectin (IVM) [a mixture of 22, 23-dihydro-avermectin B1a (80%) and 22, 23-dihydro-avermectin B1b (20%)], is a macrocyclic lactone, discovered in 1975 by Satoshi Omura as a fermentation product of the actinomycete *Streptomyces avermitilis* and developed in the early eighties to treat parasitic diseases. Its high lipophilicity and efficacy allow its use through different routes, being effective to control endo and ecto-parasites in animals and humans. Shortly after its introduction in the veterinary market, the drug was approved for human use. Nowadays, after decades of intensive, safe and effective use, IVM is indicated to treat several neglected tropical diseases, including onchocerciasis, helminthiases, and scabies. It had also been evaluated for its potential to reduce the rate of malaria transmission by killing mosquitoes (1). Overall, IVM has been widely used, demonstrating an excellent safety profile. Additionally, in the last few years, new knowledge guided the repurposing of the drug towards the treatment of other diseases. IVM antibacterial (2), antiviral^3^ and antimitotic activities (4, 5, 6) have been experimentally observed.

IVM antiviral activity against Dengue virus (3), West Nile virus (7), Venezuelan Equine Encephalitis virus (8), and Influenza virus (9), has been reported. Recently, Caly et al. (10) reported that IVM inhibits *in vitro* the replication of SARS-CoV-2 (severe acute respiratory syndrome coronavirus) using high concentrations in the range of 2.5-5 µM. Furthermore, there is now available information from a randomized clinical trial on IVM antiviral activity in SARS-CoV-2 infected patients (11). The mechanism by which IVM inhibits SARS-COV-2, seems to be the same described for other RNA viruses, i.e. inhibition of transport across the nuclear membrane mediated by importin α/β1 heterodimer, carrier of some viral molecules indispensable for the replication process (12, 13).

SARS-CoV-2 is the etiological agent of Covid-19 (coronavirus disease 2019), a viral disease causing a pandemic since December 2019, inducing from asymptomatic to life-threatening disease. It is highly transmissible with a primary respiratory entrance and airborne transmission, which explains its extensive distribution worldwide. The information available to date indicates that SARS-CoV-2 colonizes the oropharynx and nasopharynx (NP), from where is transmitted even before the appearance of any symptoms. With viral replication in this area (14), the first symptoms (odynophagia, anosmia, dry cough, and fever) and lung parenchyma colonization appear. In the context of the current COVID-19 pandemic, it is relevant to determine the best way to administer IVM to optimise its potential in vivo therapeutic usefulness.

The IVM pharmacokinetic features, based on high lipophilicity and a large volume of distribution, allow its high availability in the respiratory tract (15, 16). Thus, considering the gateway of the virus, the administration of a nasal IVM spray (N-IVM spray) intended to deposit the drug in the upper respiratory tract, could represent a practical tool to expose the of SARS-CoV-2 virus (or the cells where the viral particles are located) to high concentrations of IVM. Hence, a reduction of the viral load at the beginning of the infection, preventing viral replication, transmission and disease aggravation might be achieved.

Only limited information on inhaled IVM in rats is available (17) and to the best of our knowledge, this is the first time that a nasal IVM spray formulation is developed and its safety and pharmacokinetic performance determined in a pig model, the most appropriate animal model to use in translational research into humans. In an attempt to achieve high IVM concentrations in tissues where entry and transmission of SARS-CoV-2 occurs (where large viral loads are found at the early stages of the infection), the main goal of the work described here was to assess the safety and pharmacokinetic performance of a novel IVM-spray formulation for intranasal administration in piglets. The IVM concentration profiles measured in plasma, NPand lung tissues after the intranasal treatment (one and two applications) were compared with those achieved in the same tissues after the oral (tablets) administration of the antiparasitic dose of 0.2 mg/kg approved for human use. The work reported here is fully supported by recently available scientific evidence on both the potential preventive effect of IVM in SARS-CoV-2 transmission (18), and the concentration-dependent IVM effect on the viral load decay rate observed in a recently completed controlled clinical trial in COVID-19 infected patients (11). This clinical trial was simultaneously performed with the work described here by the same authors as a part of large public-private joint research collaboration in Argentina.

## Material and methods

### Study formulations

N-IVM spray formulation, N-IVM-free methylene blue coloured spray formulation, IVM tablets 2.0 mg and IVM tablets 0.5 mg were developed, manufactured and quality controlled according to Good Manufacturing Practices and supplied byLaboratorio Elea-Phoenix, Argentina. The N-IVM spray was designed to deliver 1 mg IVM, in a 0.1 mL puff. Each container provides 100 puffs and calibrated microdroplets to produce a high NP tissue deposit.

### Experimental animals

Forty healthy Landrace-Duroc Jersey‐Yorkshire crossbred piglets (weighing 10 to 12 kg) were used. The animals were housed in the farm of origin. They were kept with the usual diet during the trial (antibiotic-free diet) and ad libitum access to water. Management and euthanasia of the animals were performed according to approved Good Veterinary Practices (19) and Principles of Animal Welfare (20). The study was fully performed in compliance with ethical, animal procedures *and* management protocols approved by the Ethics Committee on Animal Welfare Policy of Biogenesis Bago, Argentina (Pol-UE 0001).

### Pre-trial

A pre-trial with the N-IVM-free coloured spray was performed in two animals to assure that the target tissue areas were properly covered after one dose application and to determine the sampling methodology for NP-tissue, based on the observation of colorant presence.

### Objectives and study design

The main goals of the work described here were: 1) to assess the safety of the N-IVM spray in single and double dose administration to healthy piglets, and 2) to determine the IVM concentration profiles in NP tissue, lung tissue and plasma, at different times after its intranasal spray (one and two applications) and oral administration.

The animal phase of the study was conducted at an intensive pig farming establishment (“El Campito”, SieteBochos S.R.L., Buenos Aires, Argentina). Clinical laboratory evaluations were performed at Microdiag Laboratory, La Plata, Argentina. The tissue histopathological evaluations, drug analysis and pharmacokinetic evaluation were performed at Centro de InvestigaciónVeterinaria de Tandil (CIVETAN), UNCPBA-CICPBA-CONICET, Tandil, Argentina.

The experimental design was based on a three-group animal phase study. Group 1 (22 animals) received one dose (2 mg, 1 puff/nostril) of N-IVM-spray, Group 2 (10 animals) received two (2 mg each, 1 puff/nostril) doses N–IVM-spray 12 h apart, and animals in Group 3 (8 animals) were treated with IVM (0.2 mg/kg) oral tablets. Animals with body weight from 10 to 12 kg were selected to allow an equivalent standard-dose treatment for one dose N-IVM administration and one dose oral (0.2 mg/kg) treatment. A summary of dosing and group design is shown in Table I.

**Table I:** Summary of the experimental design for the study animal phase.

|  | Group 1 | Group 2 | Group 3 |
| --- | --- | --- | --- |
| Formulation | N-IVM spray | N-IVM spray | Oral tablets |
| Treatment | 1 dose<br>(1 puff/nostril) | 2 doses<br>(1 puff/nostril)<br>12h apart | 1 oral dose |
| Dose | 2 mg | 4 mg | 0.2 mg/kg<br>(approx. 2 mg) |
| Animals (n) | 22 | 10 | 8 |
| Purpose | Safety<br>Pharmacokinetics | Safety<br>Pharmacokinetics | Pharmacokinetics |

### Safety assessment of the N-IVM spray formulation

The overall safety and local tissue tolerability of theN-IVM spray formulation were assessed. The IVM-treated animals were monitored by a careful clinical examination. Vital signs, haematological/serum biochemistry analysis and histopathology of tissues at the drug application area were assessed. All the experimental animals were carefully monitored for adverse effects throughout each dosing period.

During the first 6 h after administration and then at 12 and 24 h, careful clinical control was performed, looking for any signs of nasal or respiratory discomfort, abnormal behavior, the appearance of the stool, feed and water consumption. Immediately after each drug administration and at 2, 6, 12 and 24 h after treatment, physical/visual examination of the application sites was performed. External and internal mouth inspection was performed to determine any possible adverse effect, which included visual observation of any possible abnormal manifestation in the NP epithelium area as a consequence of the N-IVM spray application.

### Clinical laboratory evaluation

Blood samples were collected at baseline (before treatment) (12 samples) and 24 h after single-dose (6 samples) and double-dose (6 samples) treatments to perform the haematologic and serum biochemical analysis. Haematology included measurements of hematocrit, red blood cell count, hemoglobin, mean corpuscular volume (MCV), mean corpuscular hemoglobin (HCM), mean-corpuscular hemoglobin concentration (MCHC) and counts of leukocytes, band neutrophils, segmented neutrophils, eosinophils, basophils, lymphocytes, monocytes and platelets. Serum concentrations of urea, creatinine, and the activities of alanine aminotransferase (ALT), aspartate aminotransferase (AST) and alkaline phosphatase (ALP) enzymes were determined.

### Blood, nasopharyngeal and lung tissues sampling for drug measurement

Four (4) animals from each of the experimental groups were randomly selected to be euthanized at each sampling time following approved animal guidelines. The head of the animals was opened following a sagittal line, the nasal septum removed and discarded, and a scrape of NP mucosa, submucosa, turbinates and soft palate were integrated to form the NP-tissue sample. After opening the chest, a portion of the upper lobe of the right lung was obtained.

Following a single dose of the N-IVM spray administration (Group 1), samples of blood, NP and lung tissues were collected at 2, 6, 12 and 24 h post-dose to measure IVM concentrations. After the double dose N-IVM spray administration (Group 2) and oral administration (Group 3), samples of blood, NP-tissue and lung were collected at 6 and 24 h post-dosing. Plasma was separated by centrifugation at 2500 rpm for 15 min. The plasma and collected tissue samples were placed into plastic tubes and frozen at −20°C until analysis byHigh Performance Liquid Chromatography (HPLC).

### Histopathological study

NP and oropharynx epithelia were carefully examined post-mortem for assessing possible macroscopic abnormalities induced by the N-IVM spray application. Samples of NP-tissue from the soft palate region were obtained at 24 h post-administration of the N-IVM spray formulation (animals receiving one or two intranasal applications) to perform the histopathological assessment.

### Analytical development. Measurement of IVM tissue concentration profiles

Concentrations of IVM in NP-tissue, lung and plasma samples were determined by HPLC with fluorescence detection following the technique previously described (15). An aliquot of plasma and tissues were homogenized and combined with moxidectin as an internal standard. Full validation of the analytical procedures used to measure IVM concentrations in the different tissues was performed. After acetonitrile-mediated chemical extraction, IVM was converted into a fluorescent molecule using N-methylimidazole and trifluoroacetic anhydride (Sigma Chemical, St Louis, MO, USA). An aliquot (100 μl) of this solution was injected directly into the HPLC system (Shimadzu Corporation, Kyoto, Japan). The determination coefficients (r^2^) of the calibration curves for the different tissues analysed ranged between 0.989 and 0.999. The mean absolute drug recovery percentages were 94% (NP-tissue), 86% (lung tissue) and 97% (plasma). The relative error values (accuracy) was in the range between 2.9% and 9.4%. The method exhibited a high degree of inter-day precision with a coefficient of variation below 7%. The limits of drug detection were 0.45 ng/g (NP-tissue), 0.19 ng/g (lung) and 0.20 ng/mL (plasma). The limits of quantification (LOQ) were 0.70 ng/g (NP-tissue), 0.30 ng/g (lung) and 0.34 ng/mL (plasma). Concentration values below the quantitation limits were not considered for the pharmacokinetic analysis.

### Pharmacokinetic and statistical analysis of the data

The IVM concentration versus time curves obtained for each tissue/fluid after each experimental treatment were fitted with the PK Solutions 2.0 (Ashland, Ohio, US) computer software. The area under the concentration-time curves (AUC) was calculated by the trapezoidal rule (21) to determine the IVM exposure (tissue availability) at each assayed tissue. The statistical analysis was performed using the Instat 3.0 software (GraphPad Software, CA, US). IVM concentrations after the different treatments were statistically compared using a non-parametric Kruskal**-**Wallis test. The data from the hematological and biochemical determinations were compared by basic statistical analysis using the Info Stat, 2016 software.

## Results

The N-IVM spray was well tolerated after either one or two applications to the Animal model. The piglets were considered clinically healthy by specialised veterinarians throughout the whole experimental trial. All had normal skin and mucous membranes color, body condition and behaviour activity. No adverse events or intolerance were evident along the whole study period in animals treated either orally or with the spray formulation once or twice.

There were no macroscopic changes at the tissue area of spray application at different examination times. Furthermore, no histopathological changes (lesions) were observed in the mucosa or submucosa of the soft palate in spray-treated animals. A mild to moderate inflammation was observed in the tonsils (both before and after spray application), which is normal in pigs because of the immunological role of the area. The serum biochemical and hematological values did not show any alteration that would lead to adverse effects. Range values for hematological and biochemical determinations before and after the administration of the N-IVM spray formulation are summarized in Table II.

**Table II.**
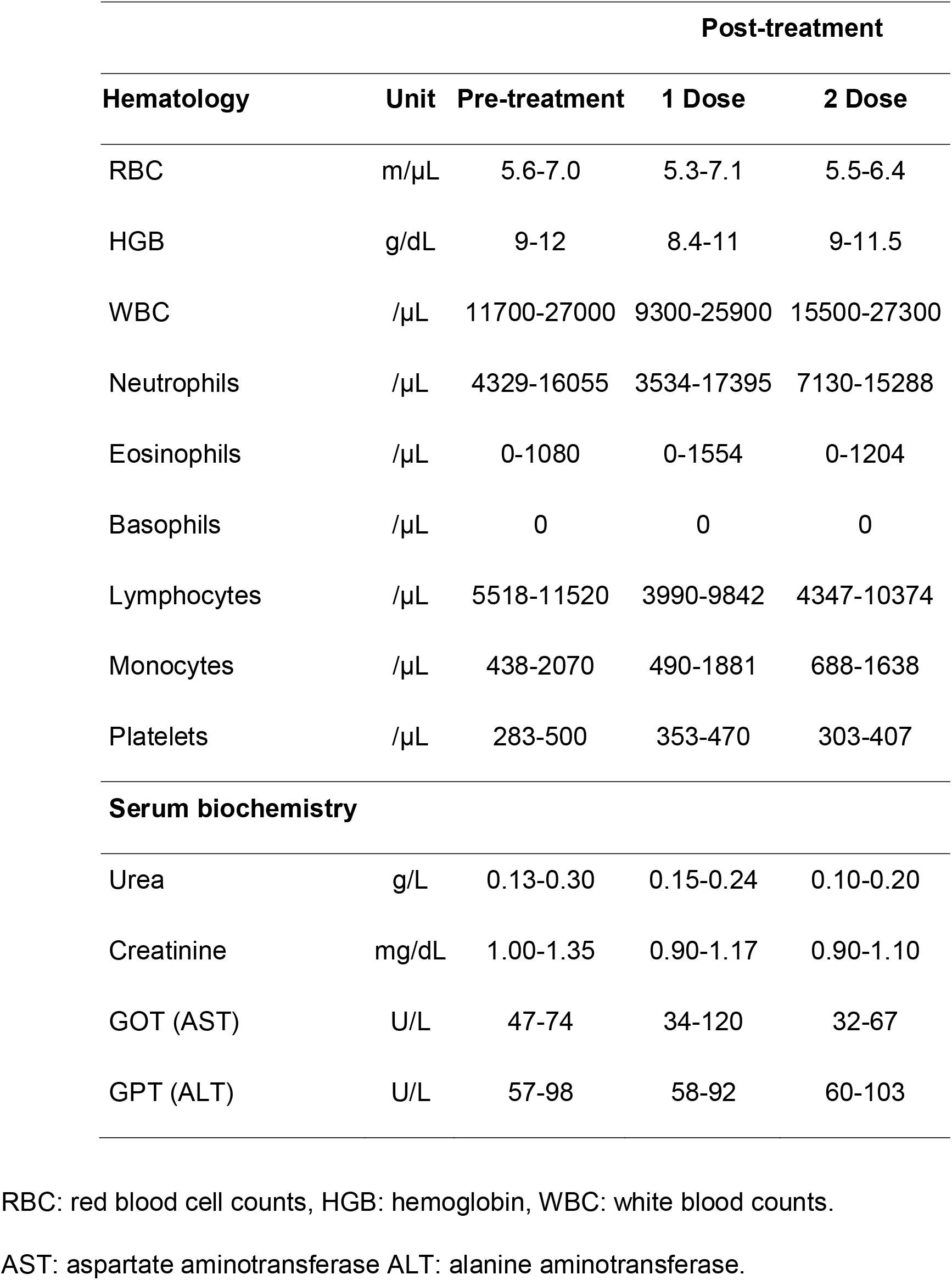
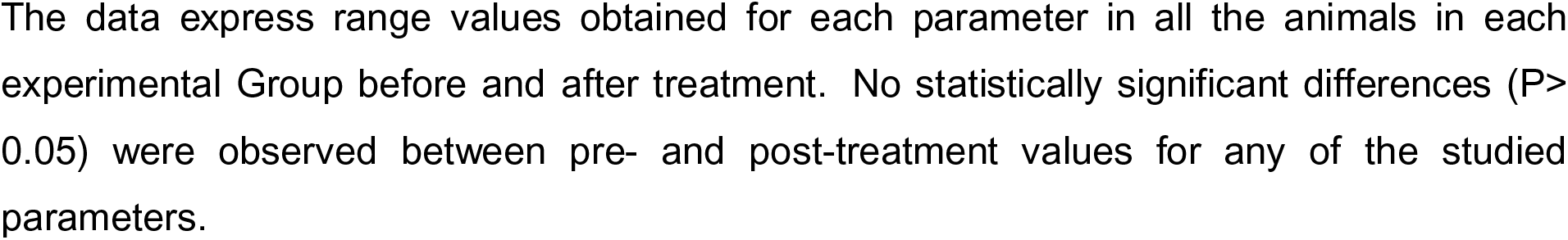
Haematological and serum biochemical range values obtained in experimental pigs before and after the treatment with either one or two doses of the N-IVM spray formulation.

IVM was recovered in plasma, NP and lung tissues following a single dose application of the N-IVM-spray formulation (Group 1). Although some degree of variability was observed in the patterns of tissue concentration among the animals treated with the spray formulation, the highest IVM concentrations were always measured in NP-tissue. Additionally, high IVM concentrations were measured in lung tissue, with a limited passage into the central compartment (systemic absorption), reflected in the low plasma levels recovered in the animals treated with N-IVM spray. The comparative IVM concentration profiles in NP-tissue, lung and plasma obtained over the first 24 h post-administration (one dose) of the N-IVM spray formulation, are shown in Figure 1.

**Figure 1.**
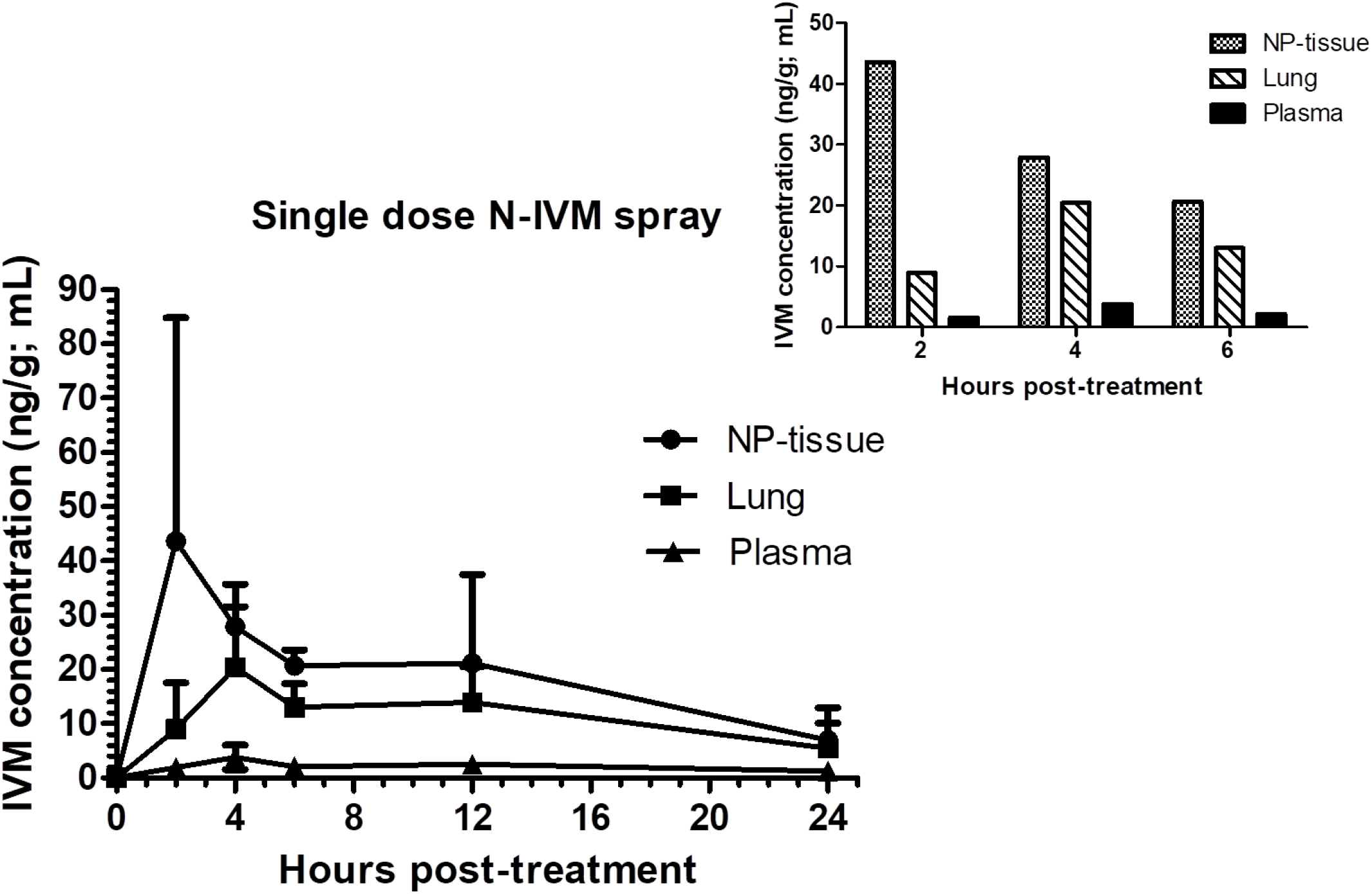
Mean ivermectin (IVM) concentration profiles in nasopharyngeal (NP) tissue, lung and plasma following the intranasal (N-IVM-spray, one dose) administration to piglets. The insert shows the comparative IVM concentrations obtained between 2 and 6 h post spray administration.

A significant positive correlation between the IVM concentrations in NP and lung tissues (r=0.735) was observed between the 4 h and 24 h post-administration. The IVM exposure (measured as AUC values) was calculated for each of the assayed tissues and plasma. The highest drug exposure after the one dose nasal application was observed at the NP and lung tissues. The drug availability (exposure) expressed as AUC values in each tissue and the relationship (ratio) between IVM exposure in lung and plasma compared to NP tissue are shown in Table III.

**Table III:** Ivermectin (IVM) exposure (availability) expressed as AUC values in nasopharyngeal (NP) tissue, lung tissue and plasma following the N-IVM-spray (one dose) administration.

|  | **IVM exposure**<br>**(mean AUC values)** | **Relationship with**<br>**NP exposure\*** |
| --- | --- | --- |
| **NP tissue** | 457ng.h/g | - |
| **Lung Tissue** | 268ng.h/g | 0.59 |
| **Plasma** | 49.5ng.h/mL | 0.11 |

\*Drug exposure (AUC) ratios in lung/NP tissues and plasma/NP tissue

IVM concentrations were also measured following the two doses of N-IVM-spray administration as well as after the treatment with the oral tablets in pigs. The repeated spray treatment increased significantly the IVM concentrations in NP and lung tissues compared to those measured after the single intranasal administration, without any significant increment on IVM concentrations in the bloodstream, reflecting a limited IVM systemic absorption. These results confirm the hypothesis of a high IVM availability in the NF tissue area following the nasal spray administration. Besides, a high distribution of IVM was observed after its oral administration, with higher lung concentration profiles compared to NP tissue measured at 6 h post-treatment. The comparative mean IVM concentrations in each tissue after the three experimental treatments are shown in Table IV.

**Table IV:** Mean (±SD) comparative ivermectin (IVM) concentrations measured in nasopharyngeal tissue, lung tissue and plasma at 6and 24 h after its intranasal (N-IVM spray) (as one and two applications) and oral administration to piglets.

| IVM concentration (ng/g; ng/mL) |  |  |  |
| --- | --- | --- | --- |
|  | N-IVM spray<br>(one 2 mg dose) | N-IVM spray<br>(two 2 mg doses, 12 h<br>apart) | Oral<br>(one 0.2 mg/kg<br>dose) |
| **Nasopharyngeal** |  |  |  |
| 6 h | 20.7 ± 2.92<sup>a</sup> | 49.1 ± 18.7<sup>b</sup> | 23.6 ± 10.8<sup>a</sup> |
| 24 h | 6.96 ± 6.01<sup>a</sup> | 10.9 ± 7.58<sup>a</sup> | 13.0 ± 5.33<sup>a</sup> |
| **Lung tissue** |  |  |  |
| 6 h | 13.0 ± 4.38<sup>a</sup> | 34.4 ± 10.2<sup>b</sup> | 54.7 ± 14.0<sup>c</sup> |
| 24 h | 5.42 ± 4.69<sup>a</sup> | 13.0 ± 9.41<sup>a</sup> | 25.4 ± 10.3<sup>b</sup> |
| **Plasma** |  |  |  |
| 6 h | 2.13 ± 0.50<sup>a</sup> | 4.95 ± 2.92<sup>ab</sup> | 8.60 ± 3.39<sup>b</sup> |
| 24 h | 1.20 ± 1.04<sup>a</sup> | 3.37 ± 2.61<sup>b</sup> | 3.65 ± 0.88<sup>b</sup> |

Different letters indicate statistically significant differences at $P < 0.05$ .

Using the oral tablet administration as a reference, the relationship between IVM concentrations recovered at 6 h in target tissues (NP and lung) after the spray application was estimated. The spray/oral concentration relationship increased significantly from 0.88 (one spray application) to 2.10 (two spray applications) in NP tissue and from 0.24 to 0.63 in lung tissue. A less marked increase for the same spray/oral ratio was observed in plasma (from 0.25 to 0.57), as it can be observed in Figure 2.

**Figure 2.**
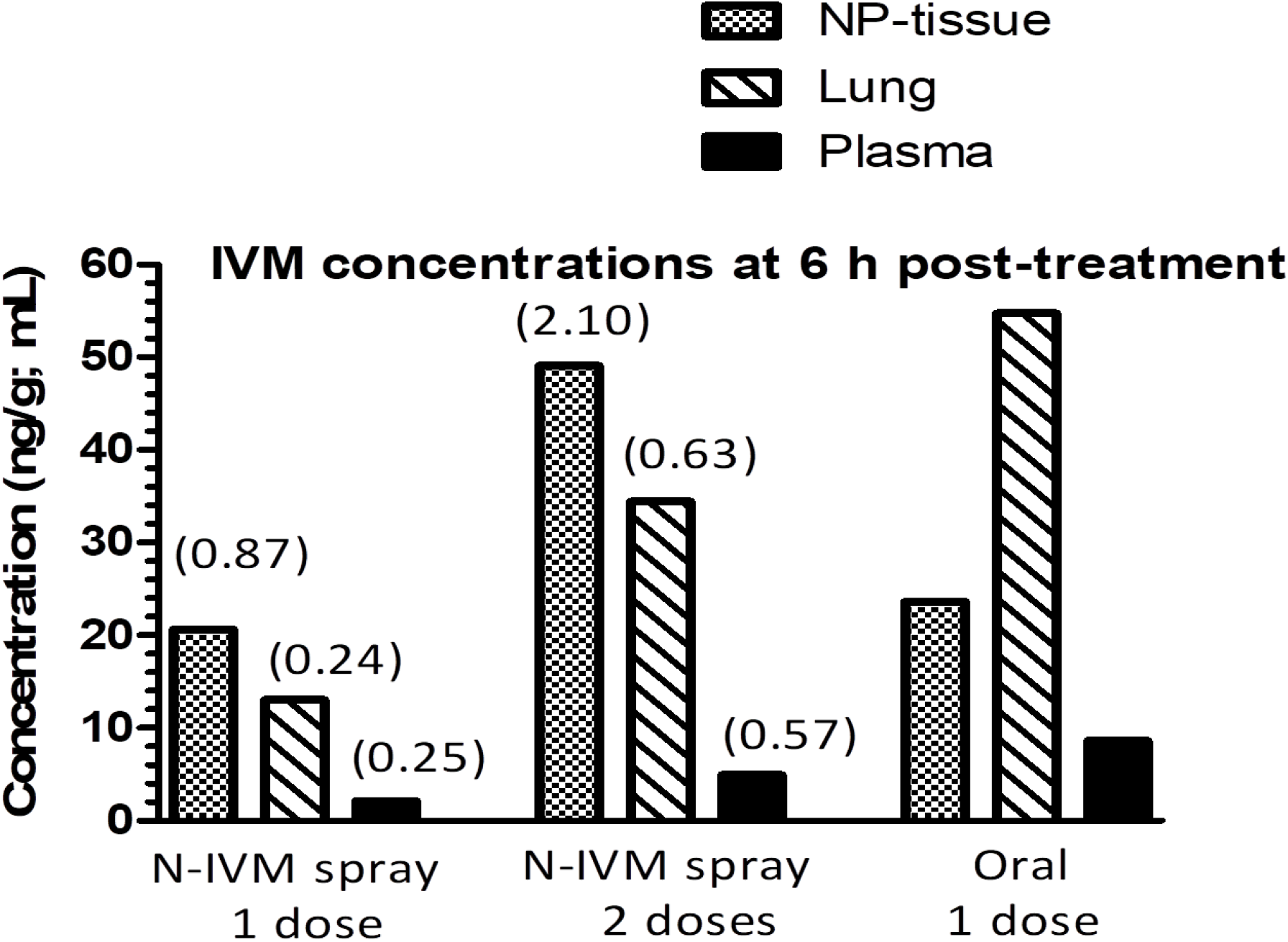
Comparative ivermectin (IVM) concentrations in nasopharyngeal (NP) tissue, lung and plasma at 6 h after oral and N-IVM-spray (one and two doses) treatment. The spray to oral IVM concentration ratios (values in brackets) are shown for each o fthe target tissues/plasma.

## Discussion

The work reported here illustrates the safety and pharmacokinetic assessment of a novel pharmaceutical formulation aimed to attain high IVM concentrations in NP and lung tissues with low systemic availability. The pig was chosen as the test animal model to assess the safety, application site tolerability and kinetic performance of the new formulation and innovative route of IVM administration. Pigs and humans have anatomical and physiological similarities. The pig is the animal species most used in translational research in studies of pathophysiology, cardiovascular and gastrointestinal surgery, preclinical toxicological testing of pharmaceuticals, and lately for the understanding of the anatomy of the respiratory system and training in lung transplantation (22).

The assessment of safety and pharmacokinetics for the novel IVM spray formulation in a pig model is described for the first time. The N-IVM-spray was shown to be safe and well tolerated. Neither clinical adverse effects, haematological, serum biochemical nor histopathological changes on the tissue area of drug application, were observed in animals treated with the spray formulation.

The work reports original data on IVM concentration profiles on NP-tissue after an intranasal and oral administration in a pig animal model. While the repetition of the intranasal dose at a 12 h interval determined a significant increase in IVM concentrations in NP and lung, both identified as target tissues for SARS-CoV-2, only minimal drug systemic exposure (low plasma levels) was observed. After 0.2 mg/kg oral tablet administration, IVM plasma concentration levels were in agreement with those previously published for piglets (23, 24). IVM was well absorbed and extensively distributed, resulting in higher concentrations in lung tissue than in plasma (see Figure 2), a pattern already observed in cattle subcutaneously treated with IVM15, and confirmed by a recent pharmacokinetic simulation report looking at IVM lung exposure in humans. Using a minimal physiological pharmacokinetic model, this work establishes that a lung mean concentration as high as 193 ng/g may be achieved after a single oral treatment of 30 mg (16).

The variability observed among treated animals on the patterns of IVM concentration in NP and lung tissues may be related to the fact that it is not possible to control the ventilator state of the animals at the moment of the spray drug application. As a consequence, some animals could have different degrees of inspiration during the spray administration, which may help to understand individual variation on drug concentrations measured both in NP and lung tissues. The situation could be different if this type of N-IVM-spray is used by humans, since the user could be instructed to slightly inspire or remain in apnea, with a predictable increase in drug penetration into the respiratory tree achieving higher lung drug concentrations.

Several randomized controlled trials are ongoing to investigate the efficacy of IVM against COVID-19, using oral treatments at different doses. Moreover, many uncontrolled oral treatments are using the approved antiparasitic dose of 0.2 mg/kg. Recently reported data18 has shown the potential preventive effect of IVM in SARS-CoV-2 transmission. Additionally, we have recently demonstrated the concentration-dependent IVM effect on the viral clearance in a controlled clinical trial in COVID-19 infected patients (11). The scientific evidence of the in vivo effects of IVM on reducing the SARS-CoV-2 viral load gives prominence to the data on the assessment of the spray formulation in a pig model described here.

It may be expected that repeated intranasal administration increases IVM concentrations in NP-tissue and lungs. Asa low systemic absorption was observed, and considering the intrinsic safety of the drug, there would be no anticipated significant risks of IVM systemic toxicity after repeated nasal administration. Compared to the oral administration, the intranasal administration in humans may provide fast, high and persistent IVM concentration at the NP tissue area at much lower doses. For instance, in a 60 kg body weight person, higher IVM concentrations in NP tissue may be reached with 4 mg of N-IVM (2 spray doses) than giving 12 mg orally (1 dose 0.2 mg/kg). Thus, the daily administration of one puff in each nostril to health workers would allow the persistence of high IVM concentrations in NP epithelium during an entire working period. The design and execution of clinical trials to confirm safety and to determine the efficacy of N-IVM spray should evaluate these potential benefits. This could include recently diagnosed COVID-19 patients, their close contacts, and/or a preventive usage in health workers. Based on the pharmacokinetic data shown here, the administration of more than one puff per nostril a day would allow IVM accumulation in NP tissue, reaching a local drug exposure not feasible to be achieved by the oral route. In the same direction, further research is also needed to evaluate the potential advantages of a combined nasal plus oral treatment regimen to further contribute to IVM repurposing in COVID 19 therapy.

## Acknowledgements

The authors wish to thank all those collaborators (at each of the involved institutions) who have anonymously contributed with the execution of the work reported here.

## Funding

The work described here was mainly supported by funding from Laboratorio Elea Phoenix, Argentina. The Consejo Nacional de InvestigacionesCientíficas y Técnicas (CONICET), Argentina, The Facultad de CienciasMédicas de la Universidad Nacional de La Plata, Argentina, and INCAM S.A. partially contributed through payment of salaries for several of the authors in this article. The funders had no role in study design, data collection and interpretation, or the decision to submit the work for publication

## Author Contributions

J. Errecalde. Protocol design, IVM spray design. Animal phase work (Spray administration and sampling). Data analysis. Overall integration/discussion of the data. Manuscript writing.

A. Lifschitz. Protocol design. HPLC analysis. PK data analysis. Overall integration/discussion of the data. Manuscript writing.

G. Vecchioli. Protocol design. Data analysis. Overall integration/discussion of the data. Manuscript writing.

L. Ceballos. Analytical development. Method validation. HPLC analysis. Data integration

F.Errecalde. Animal phase work (treatments/sampling).

M. Ballent. Analytical development. Method validation.

G. Marín. Dosage calculation. Manuscript’s revision.

M. Daniele. Animal phase work (treatments/sampling).

E. Spitzer, F. Toneguzzo, S. Gold., Protocol Design. Pharmaceutical Spray development. Regulatory Discussion. Overall Integration/analysis/discussion of the data.

A. Krolewiecki. Protocol design. Overall integration/discussion of the data.

L. Alvarez. Protocol design. Overall discussion of the data. Manuscript writing.

C. Lanusse. Protocol design. Overall integration/discussion of the data. Manuscript writing.

## Notes

### Competing Interest Statement

The authors have declared no competing interest.

